# Structural asymmetry along protein sequences and co-translational folding

**DOI:** 10.1101/2020.09.26.314609

**Authors:** John M. McBride, Tsvi Tlusty

## Abstract

Proteins are translated from the N- to the C-terminal, raising the basic question of how this innate directionality affects their evolution. To explore this question, we analyze 16,200 structures from the protein data bank (PDB). We find remarkable enrichment of *α*-helices at the C terminal and *β*-sheets at the N terminal. Furthermore, this *α*-*β* asymmetry correlates with sequence length and contact order, both determinants of folding rate, hinting at possible links to co-translational folding (CTF). Hence, we propose the ‘slowest-first’ scheme, whereby protein sequences evolved structural asymmetry to accelerate CTF: the slowest-folding elements (*e*.*g. β*-sheets) are positioned near the N terminal so they have more time to fold during translation. Our model predicts that CTF can be accelerated, up to double the rate, when folding time is commensurate with translation time; analysis of the PDB reveals that structural asymmetry is indeed maximal in this regime. This correspondence is greater in prokaryotes, which generally require faster protein production. Altogether, this indicates that accelerating CTF is a substantial evolutionary force whose interplay with stability and functionality is encoded in sequence asymmetry.

---

All proteins are translated sequentially from the N- to the C-terminal, and are thus inherently asymmetric [1]. One example of such N-to-C asymmetry is signal peptides, which enable translocation across membranes, and are located at the N terminal [2]. This raises the question of whether and how the universal unidirectionality of protein production is leveraged to gain evolutionary advantage. Here we examine structural data from the protein data bank (PDB) in search of traces of such adaptation. We analyzed the distribution of secondary structure along the sequence for 16,200 PDB proteins, finding two striking patterns of asymmetry. First, disordered residues are principally located at the ends of sequences, and depleted towards the middle. Second, *β*-sheets are enriched by 55 % near the N terminal, while *α*-helices are enriched by 22 % at the C terminal. This *α*-*β* asymmetry peaks at intermediate values of sequence length and contact order – which both correlate negatively with folding rate – indicating a possible link between secondary structure asymmetry and folding.

Hence, we further explore the possibility that *α*-*β* asymmetry may accelerate protein production, and is therefore a signature of evolutionary adaptation. Production of functional proteins from mRNA comprises two concerted processes: translation and folding. The rate of translation is limited by trade-offs between speed, accuracy and dissipation [3–6]. Folding quickly has certain advantages: unfolded proteins lead to aggregation, putting a significant burden on the cell [7–9]; faster folding allows quicker responses to environmental changes [10, 11]. Moreover, organisms whose fitness depends on fast self-reproduction would benefit from accelerated protein production that can shorten division time [12, 13]. Proteins begin folding during translation [14–19]. Thus, in principle, faster production times may be achieved if proteins finish folding and translation at around the same time. This co-translational folding (CTF) enables adaptations that increase yield and kinetics of protein production [17–21]. For example, nascent peptides interact with ribosomes and chaperones to reduce aggregation and misfolding [22–26], while translation rates can be tuned to facilitate correct folding [27–30]. Specifically, we ask if structural asymmetry may have evolved for fast and efficient production via CTF.

We show that the structural asymmetry observed in proteins is consistent with a scheme for accelerating CTF based on the sequential nature of translation and the heterogeneity of folding rates along the sequence [31, 32] – *e*.*g. β*-sheets fold much slower than *α*-helices [33]. In the proposed *slowest-first* scheme, protein sequences take advantage of this heterogeneity by evolving structural asymmetry: the slowest-folding structures are enriched at at the N-terminal [34–39], so that they are translated first and have more time to fold. A simple model predicts that, under this scheme, production rate can be almost doubled when folding time is equivalent to translation time. To examine this hypothesis, we estimate the ratio of folding to translation time of the PDB proteins and compare it with their *α*-*β* asymmetry, finding that asymmetry peaks when folding time is commensurate with translation time. In this region, proteins are twice as likely to exhibit *α*-*β* asymmetry that favours the slowest-first scheme. We see more evidence for this scheme in prokaryotic proteins, which is consistent with prokaryotes’ greater need for fast protein production due to more frequent cell division. Taken together, these findings suggest that proteins sequences have been adapted for accelerated CTF via structural asymmetry.

## Results

### Protein secondary structure is asymmetric

Given the vectorial nature of protein translation, one may expect corresponding asymmetries in protein structure. To probe this, we study a non-redundant set of 16,200 proteins from the Protein Data Bank (PDB) [40]. We find that these PDB proteins exhibit significant asymmetry in secondary structure (Fig. 1A-B): the first 20 residues at the N terminal are 55 % more likely to form sheets, and the first 20 residues at the C terminal are 22 % more likely to form helices. This asymmetry is stronger for prokaryotic proteins (72 %; 20 %) than for eukaryotic proteins (20 %; 28 %). The substantial *α*-*β* asymmetry points to an evolutionary driving force which we further investigate.

**Figure 1:**
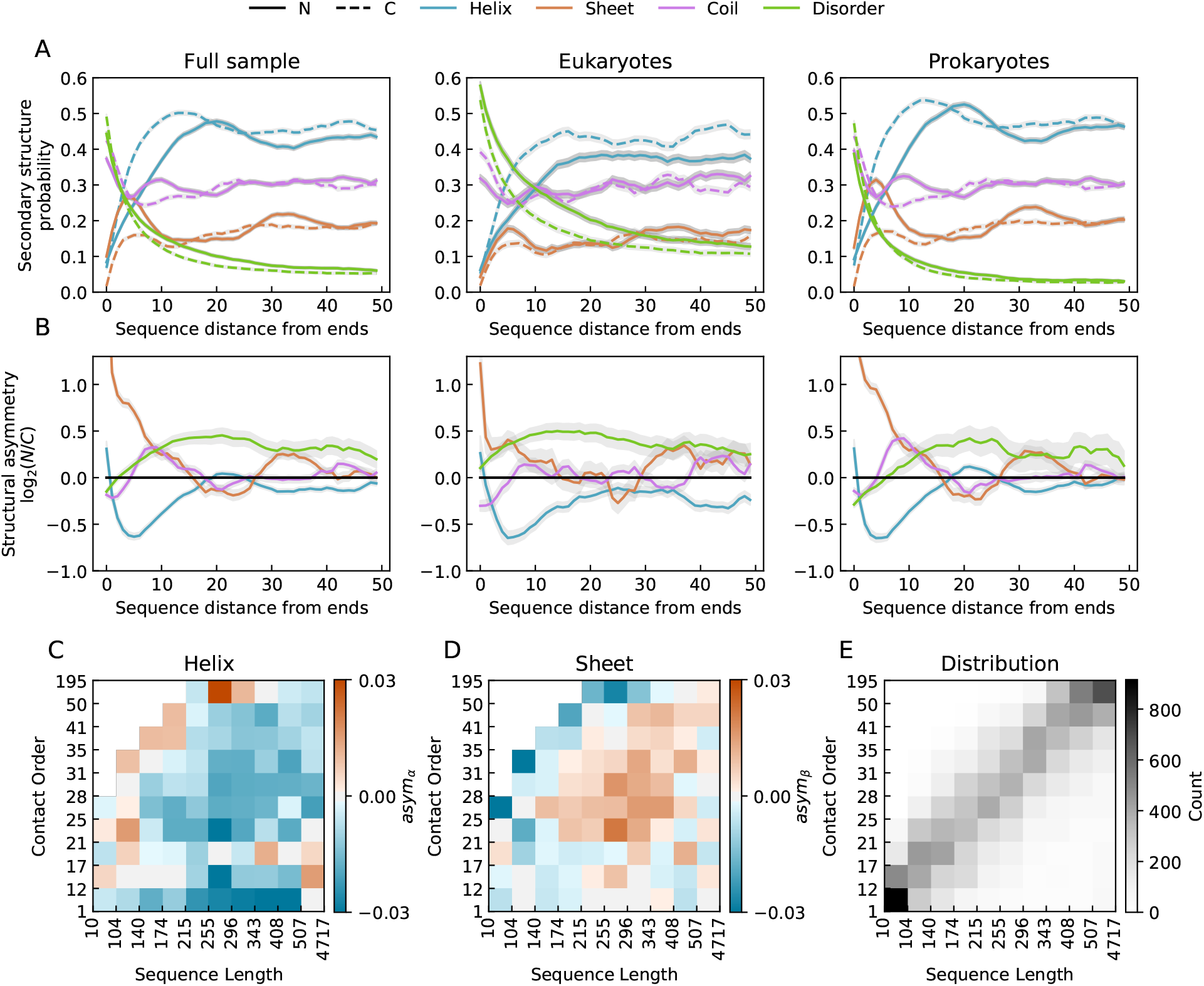
(A) Distribution of secondary structure along the sequence as a function of distance from the N- and C-terminal, and (B) the structural asymmetry – the ratio of the N and C distributions (in log_2_ scale; ± 1 are 2:1 and 1:2 N/C ratios) – for all 16,200 proteins (left), 4,702 eukaryotic proteins (middle), and 10,966 prokaryotic proteins (right). Shading indicates bootstrapped 95% confidence intervals. C-D: Mean asym_*α*_ (C) and asym_*β*_ (D) as a function of sequence length and contact order (Eq. 1). The data is split into deciles and the bin edges are indicated on the axes. E: Distribution of proteins according to contact order and sequence length.

In both N- and C-terminals, the *α*-helix and *β*-sheet distributions exhibit *periodicity* in the positioning of these elements along the sequence. This periodicity is matched by several *αβ*-type protein folds where *α*-helices and *β*-sheets are arranged in alternating order (SI Fig. 1). These folds tend to be more abundant in prokaryotic proteins (SI Table 1); for example, ferredoxin-like folds exhibit high *α*-*β* asymmetry, significant periodicity at the N-terminal, and are *∼*3 times more common in prokaryotes.

**Table 1:** The probability of that a prokaryotic protein has a specific fold, *P*_Prok_, compared with the probability of that a eukaryotic protein has a specific fold, *P*_Euk_. Results are shown for the 9 most common SCOP folds [3] identified in PDB data used in this study.

| SCOP Class | SCOP ID | Count | Description | $P_{\text{Prok}}$ | $P_{\text{Euk}}$ | Probability ratio |
| --- | --- | --- | --- | --- | --- | --- |
| a/b | 2000031 | 171 | TIM beta/alpha-barrel | 0.06 | 0.03 | 1.94 |
| a/b | 2000148 | 171 | SDR-type extended Rossmann fold | 0.06 | 0.04 | 1.33 |
| a+b | 2000014 | 115 | Ferredoxin-like | 0.04 | 0.01 | 2.85 |
| a/b | 2000088 | 76 | Methyltransferase-like | 0.02 | 0.02 | 1.10 |
| a+b | 2000303 | 65 | Acyl-CoA N-acyltransferases (Nat) | 0.02 | 0.01 | 4.14 |
| a | 2000002 | 62 | Globin-like | 0.00 | 0.07 | 0.07 |
| a/b | 2000016 | 58 | Rossmann(2x3)oid (Flavodoxin-like) | 0.02 | 0.01 | 1.98 |
| a/b | 2000607 | 58 | PLP-dependent transferase-like | 0.02 | 0.02 | 1.04 |
| a/b | 2000027 | 56 | ClpP-type beta-alpha superhelix | 0.02 | 0.00 | nan |

### Disordered regions are more abundant at the ends

The distribution of disordered regions exhibits a different pattern of asymmetry: disordered residues are enriched at both ends of proteins compared to the middle [41, 42]. Eukaryotic proteins are significantly more disordered, where the probability of disorder is well approximated by *∼D*^*−*0.5^, where *D* is the distance from the end, while in prokaryotic proteins the probability of disorder decays as *∼ D*^*−*1^. Proteins also tend to be more disordered at the N terminals [41]: eukaryotic proteins are 30 % more likely to be disordered within the first 100 residues of the N terminal compared to the C terminal (prokaryotes: 17 %). Although prokaryotic proteins are less disordered than eukaryotic ones, the ratio of the numbers of residues in *β* sheets and *α* helices is the same.

### Structural asymmetry correlates with sequence length and contact order

To better understand the *α*-*β* asymmetry, we examined correlations with sequence length, *L*, and contact order, CO. CO is the average sequence distance between intra-protein contacts,

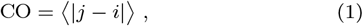

where *i* and *j* are pairs of residue indices for each contact [43]. High CO means that native contacts require large-scale movements to form, thus increasing folding time.

To quantify secondary structure asymmetry, we calculate the magnitude of asymmetry normalized by length,

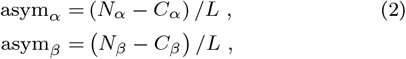

where *N*_*α*_ (*N*_*β*_) and *C*_*α*_ (*C*_*β*_) are the number of residues in a *α*-helices (*β*-sheets) in the N and C halves of a protein sequence.

We find that *α*-*β* asymmetry is a non-monotonic function of both *L* and CO (Fig. 1C-D). In particular, there is a region of intermediate length (174 *−* 408) and intermediate CO (21 *−* 35) where structural asymmetry is most apparent. The fact that both quantities correlate negatively with folding rate (*L, r* = *−*0.68; CO, *r* = *−*0.64; SI Fig. 2) [43–45], taken together with proteins’ inherent asymmetry due to vectorial translation, leads us to suspect that the origins of this *α*-*β* asymmetry may be related to co-translational folding.

### Co-translational folding appears to be widespread

During protein production, the ribosome advances along the mRNA from the N to the C terminal (Fig. 2A). Each mRNA segment encoding a structural element (red dashed segment), in turn, enters the ribosome where it is translated and passes through the ribosome channel. The time it takes this segment to clear the *∼*10 nm long ribosome tunnel, *τ*_ribo_, is the sum of the segment’s translation time *τ*_seg_ and the potential delay *τ*_delay_ until the onset of co-translational folding (CTF) once the segment exits the ribosome and is free of steric constraints [17, 46, 47].

**Figure 2:**
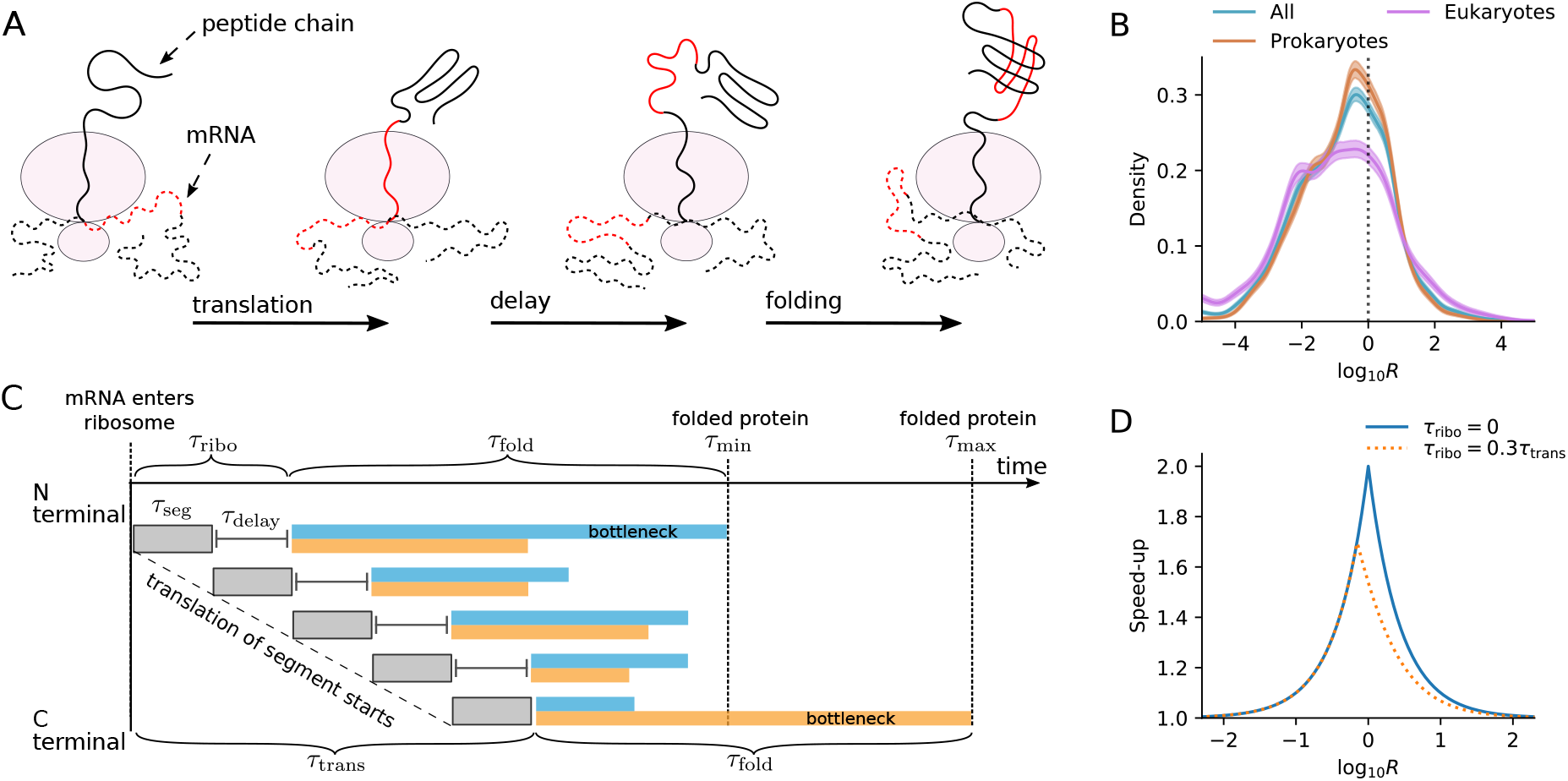
Co-translational folding and the slowest-first mechanism. A section of mRNA (red) is translated to a protein segment (left), which translocates through the ribosome channel (middle), and undergoes folding once the full segment has been translated and is free from steric constraints (right). B: Distribution of *R* = *τ*_fold_*/τ*_trans_, the ratio of folding to translation time (Eq. 3), for our entire sample, prokaryotic proteins, and eukaryotic proteins. Solid lines are kernel density estimation fits to histograms; dotted line indicates *R* = 1; shading indicates bootstrapped 95 % confidence intervals. C: Production timeline: Proteins gain functionality after translation and folding, proceeding from the N-terminal (top) to the C-terminal (bottom), where each segment begins folding after translation (*τ*_seg_) with some delay (*τ*_delay_). Two sequences (blue/orange blocks) consist of a set of structural elements with the same folding times (blocks of equal length). Production time is shortened in the blue sequence by asymmetric ordering of the folding time along the protein sequence: slow-folding sections are at the N terminal and fast-folding sections at the C terminal. Blocks for *τ*_seg_ and *τ*_delay_ are drawn the same length for simplicity. D: Theoretical maximum speedup of production rate as a function of *R* and *τ*_ribo_ (Eq. 4).

In principle, one way to maximise the rate of production and to minimise aggregation is by making proteins fold faster than they are translated, or at a similar rate. We can obtain a rough approximation of how often this occurs by estimating folding rates and translation rates of proteins. We estimate the folding rate *k*_fold_ using a power law scaling with length fitted to data from the protein folding kinetics database (PFDB) [45] (Methods). We assume an average translation rate *k*_trans_ that depends on the organism. Thus we can estimate the ratio *R* of folding time *τ*_fold_ to translation time *τ*_trans_,

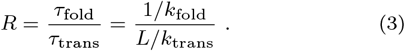

The estimated *R* distribution exhibits a peak in the region of commensurate time *R ≈* 1 (Fig. 2B). For the 68 % of proteins (CI 53 *−* 88 %, SI Fig. 3) that lie in the region *R ≤* 1, folding may be quicker than translation, indicating that CTF is common. In comparison, a more rigorous method estimated that in 37 % of proteins in *E. coli*, at least one domain will fully fold before translation finishes [48]. Examining prokaryotic proteins and eukaryotic proteins separately reveals a sharper peak in the *R* distribution for prokaryotic proteins in the region of commensurate folding and translation times, 1*/*10 *< R <* 10. Notably, a greater fraction of prokaryotic proteins (56 %) are in this regime compared to eukaryotic proteins (41 %).

### Folding rate asymmetry can speed up co-translational folding

Fig. 2C shows the production timeline of a protein whose folding time *τ*_fold_ is determined by a rate-limiting fold [32, 29–51]. This bottleneck may represent slow kinetics in secondary/tertiary structure formation, or formation of misfolded intermediates. If the bottleneck is located at the N terminal (blue blocks in Fig. 2C), then the production time is minimal, *τ*_min_ = max(*τ*_fold_ + *τ*_ribo_, *τ*_trans_). In the other extreme, if the rate-limiting fold is located at the C terminal (orange blocks in Fig. 2C), production time is maximized [52], *τ*_max_ = *τ*_fold_ + *τ*_trans_. In this case, the last element can escape the ribosome quickly after being translated (*τ*_delay_ *≈* 0) since it is not delayed by downstream translation [53]. Thus, production rate can be accelerated by a factor,

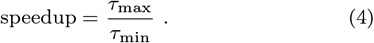

In the limit *τ*_ribo_ ≪ *τ*_trans_, one finds from Eqs. 3-4 that the speedup as a function of *R* = *τ*_fold_*/τ*_trans_ (Fig. 2D) is

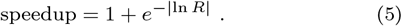

A maximal, twofold speedup is achieved when translation time equals folding time, *R* = 1, and taking *τ*_ribo_ *>* 0 shifts this maximum towards *R <* 1.

### Structural asymmetry is maximum for commensurate folding and translation times

The speedup curve (Fig. 2D) implies that proteins can benefit the most from structural asymmetry when *R* = *τ*_fold_*/τ*_trans_ *≈* 1. Hence, we estimate the magnitude of *α*-*β* asymmetry as a function of *R*, and plot the distributions in Fig. 3A. At intermediate *R*, the means of the distributions shift away from zero, indicative of strong bias.

To capture the magnitude of these shifts we calculate the N terminal enrichment, *E*, defined as the fraction of proteins with positive asymmetry (*i*.*e*. enriched at the N terminal) minus the fraction of proteins with negative asymmetry (enriched at the C terminal), for both helices and sheets:

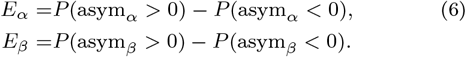

Fig. 3B shows that in the *R* decile with maximum asymmetry, proteins in the PDB are 2:0 times as likely to be enriched in *β*-sheets in the N terminal, while *α*-helices are 1.9 times more likely to be found in the C-terminal half. This maximum is found when *−*0.6 *≤* log_10_ *R ≤* 0.1 (the 95 % confidence intervals for the *k*_fold_ estimate give *−*1.1 *≤* log_10_ *R ≤* 0.7; SI Fig. 4). This region of maximal asymmetry overlaps with the region of maximal speedup (Fig. 2D, Eq. 5), suggesting that asymmetry evolves because it enhances CTF.

### Prokaryotes exhibit greater asymmetry than Eukaryotes

We looked at *α*-*β* asymmetry for prokaryotic and eukaryotic proteins separately, finding that when asymmetry is maximum, prokaryotes exhibit more asymmetry than eukaryotes – sheets are 36 % more likely to be enriched at the N terminal 218 in prokaryotes compared to eukaryotes (Fig. 3C). Typically, prokaryotic cells divide more frequently than eukaryotic cells [13], and thus have a greater need for fast production of functional proteins. The analysis is therefore consistent with the slowest-first scheme that implies that the stronger pressure on prokaryotes should lead to greater asymmetry.

**Figure 3:**
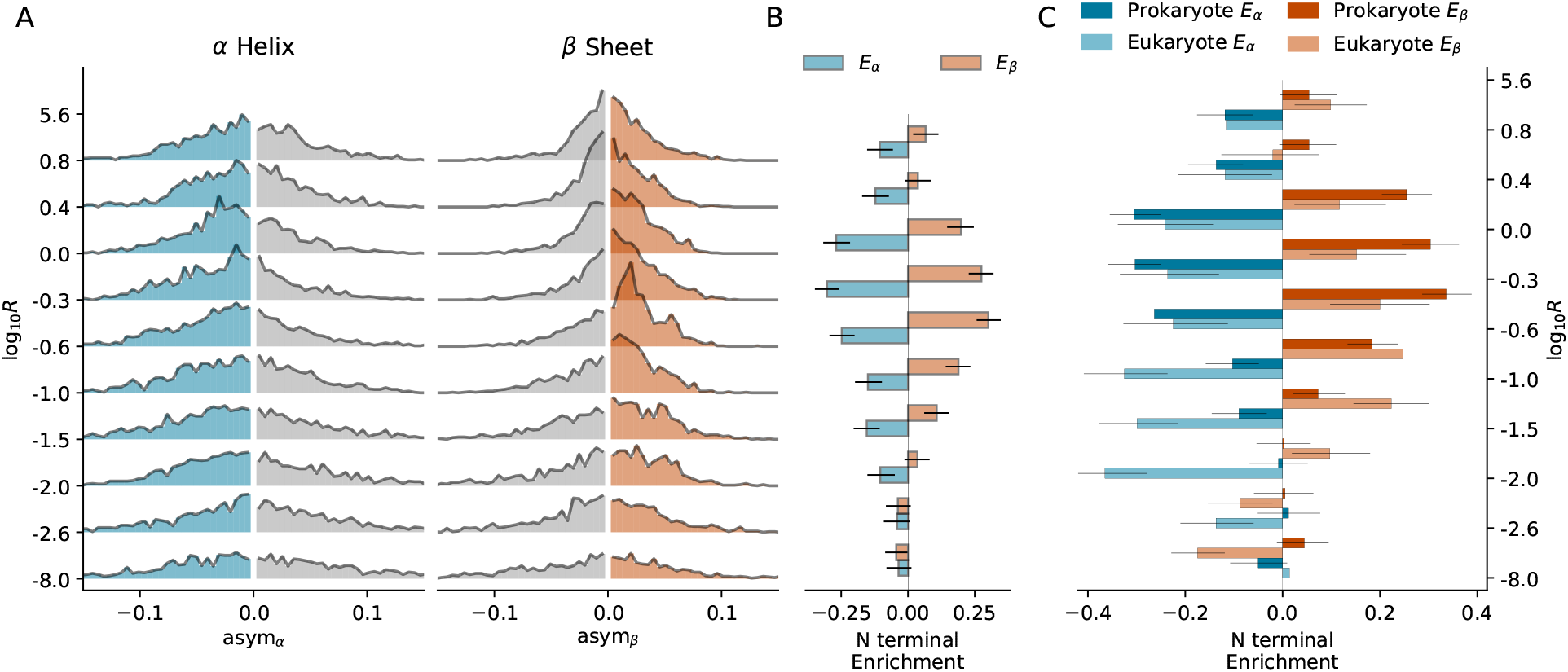
A: *α*-*β* asymmetry distributions as a function of *R*, the folding/translation time ratio (Eq. 3). Proteins are divided into deciles according to *R*; bin edges are shown on the y-axis. B: N terminal enrichment – the degree to which sheets/helices are enriched in the N over the C terminal (Eq. 6) – is shown for the deciles given in B. C: N terminal enrichment as a function of *R* for 4,702 eukaryotic proteins and 10,966 prokaryotic proteins. Proteins are divided into bins according to *R*; bin edges, shown on the x-axis, are the same as in A-B. Whiskers indicate bootstrapped 95% confidence intervals.

### Multi-domain proteins are optimized for CTF via distinct mechanisms

Multi-domain proteins can be potentially adapted at two levels: within domains, and between domains (Fig. 4A). To test this, we isolated individual domains in the PDB (using Pfam) [54], and calculated CO and *α*-*β* asymmetry for each domain as in Fig. 3. While intra-domain optimization of secondary structure clearly occurs within single-domain proteins, it is much weaker within multi-domain proteins (Fig. 4B-C). Inter-domain optimization entails ordering the slowest-folding domains at the N terminal, for which we find no significant bias (SI Fig. 5). Instead, we find that as the number of domains increases, the CO of individual domains decreases (Fig. 4D). Thus CTF is maintained in multi-domain proteins mostly by using faster-folding domains throughout.

**Figure 4:**
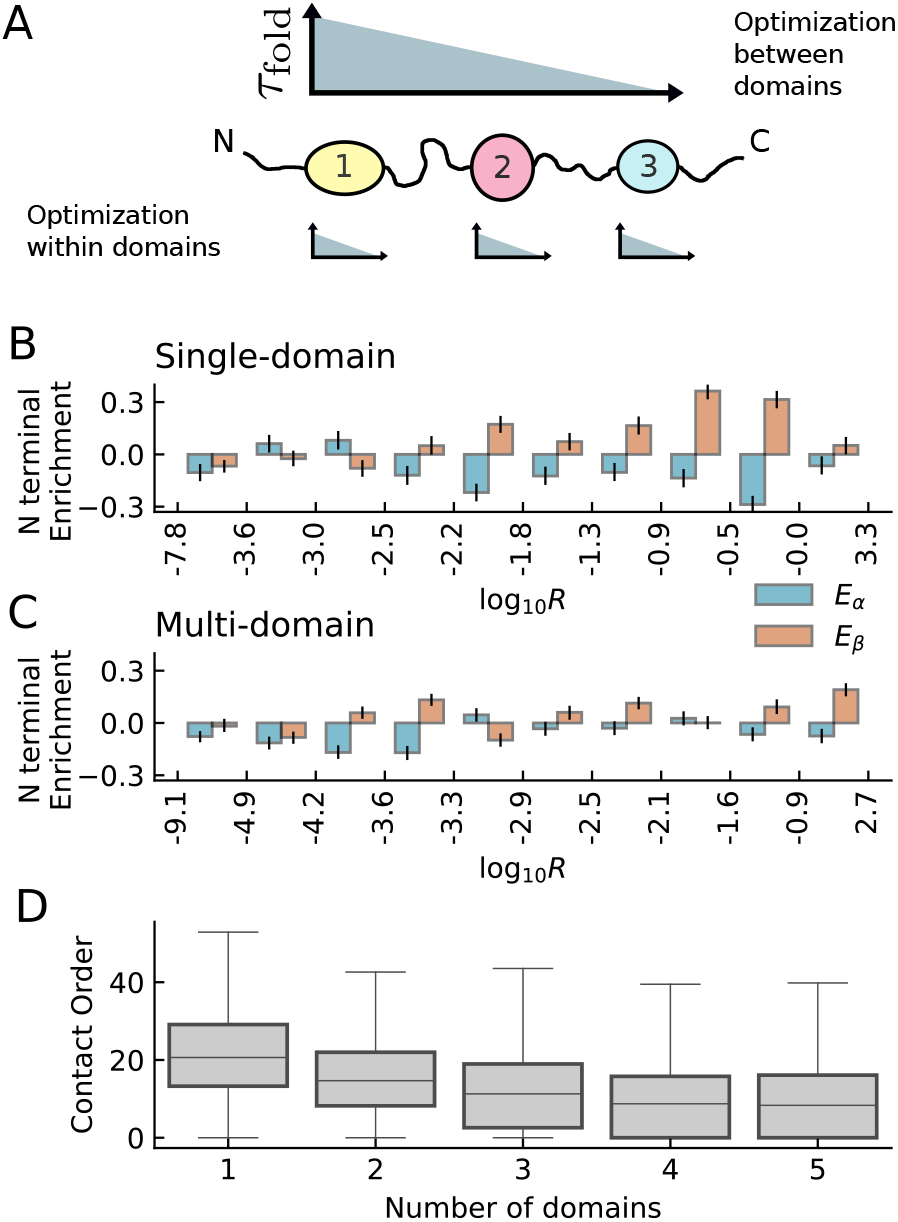
A: Multi-domain proteins can be optimized via asymmetry between domains, and/or within domains. B-C: N terminal enrichment within domains as a function of *R* for single-domain proteins (B: 14,442 domains) and multi-domain proteins (C: 23,832 domains). Domains are split into deciles based on *R*, and the bin edges are shown on the x-axis; whiskers indicate bootstrapped 95% confidence intervals. D: Domain contact order distributions for proteins with different numbers of domains.

## Discussion

### Selection pressures vary

We examined the hypothesis that proteins are selected for CTF to hasten protein production and reduce aggregation/misfolding, but this may not be equally true for all proteins. As an example, we showed that in prokaryotes, which have a greater burden of cell growth, proteins tend to have more asymmetry than in eukaryotes. More generally, CTF may be hindered in some proteins by interactions with the ribosome [55]. Long-lived proteins [56] may derive little benefit from an increase in production speed. On the other hand, proteins produced in large quantities need to fold quickly as aggregation can increase non-linearly with concentration [57]. These predictions can be tested when sufficient data for protein lifespan [58], expression levels [59], and structure become available. While we showed that *α*-*β* asymmetry is apparent in a broad set of proteins, further analysis of an extended data set may be able to detect the sub-classes of proteins that will benefit most from *α*-*β* asymmetry.

### CTF for multi-domain proteins is more complex

Multi-domain proteins exhibit less asymmetry than single-domain proteins. Due to interactions between domains [26, 29, 46, 47, 60], optimization via asymmetry may not be feasible — instead, a safe strategy is to fold each domain before translating subsequent domains. To explain the lack of intra-domain *α*-*β* asymmetry (Fig. 4C), we propose a simple mechanical argument. When a *β*-sheet forms, the protein chain contracts. This results in a pulling force on both the ribosome [61, 62], and on any upstream domains. This extra resistance to *β*-sheet formation may preclude the early formation of *β*-sheets at the N terminal side of a domain. If this is true, then the domain in position 1 should still exhibit *α*-*β* asymmetry; we currently lack sufficient statistical power to conclusively test this (SI Fig. 6). Further tests could look at CTF of a *β*-rich domain in the the presence or absence of an upstream domain [63, 64].

### Suggested experiments for circular permutants

To experimentally test the slowest-first mechanism, we suggest studying CTF of multiple proteins with *R ≈* 1, which differ in asym_*α*_ and asym_*β*_. In particular, we propose to use proteins whose sequences are related by *circular permutation*, while having identical structures [65–68]. Circular permutants with opposite structural asymmetry, as the example in Fig. 5, should fold at significantly different rates. Additional experimental control of *R* is possible via synonymous codon mutations [69] or *in vitro* expression systems [16]. Thus, one can test whether asymmetry in secondary structure can lead to acceleration of CTF, and how this depends on *R*.

**Figure 5:**
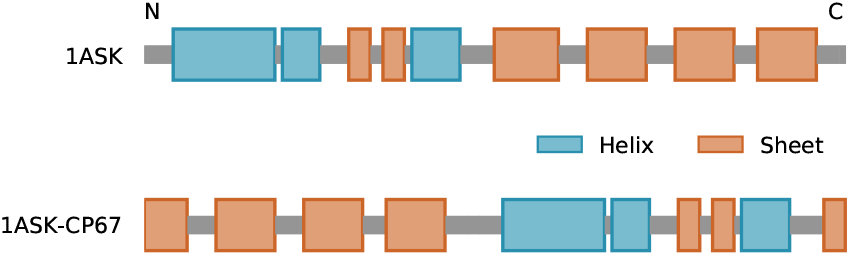
Secondary structure for nuclear transport factor 2 H66A mutant (PDB: 1ASK [70]) and a circular permutant, 1ASK-CP67, which may fold faster during translation.

### Disorder is enriched at both sequence ends

The N and C terminals principally share a notable tendency for disorder near the end, which suggests that they are affected by the same physical *end effect*. The amino acid at the end is linked to the chain by only one peptide bond, leaving it more configurational freedom than an amino acid in the centre of the protein, which is constrained by two bonds. This entropic contribution to the free energy of the loose ends, of order *k*_*B*_*T*, can induce disorder in marginally stable structures.

Since disordered regions do not need time to fold, placing them towards the C-terminal gives the other residues more time to fold. Yet, we find a similar, slightly stronger, tendency for disorder near the N-terminal (green curves in Fig. 1B), particularly in eukaryotes. This may result from other determinants of protein evolution; *e*.*g*., disordered regions tend to interact with some ribosome-associating chaperones [71, 72]. If disorder at the N terminal is related to chaperones, we expect that asymmetry will be higher for slow-folding proteins as they are more prone to aggregation. We find that bias for disorder at the N-terminal is strongest for slow-folding proteins (high *R, L* and CO; SI Fig. 7), but only for prokaryotes, not eukaryotes. Given the absence of a correlation between *R, L* and CO and disorder asymmetry in eukaryotic proteins, the question of why eukaryotic proteins are more disordered at the N terminal remains open.

### Considering tertiary structure

We used secondary structure as a proxy for folding rate, but there are also contributions from tertiary structure. To test this assumption, we ran coarse-grained simulations of CTF of three structurally-asymmetric proteins while varying *R*, for both the original sequence and of the reverse sequence. We find that these proteins fold faster when *β*-sheets are translated first, in the relevant region of *R ∼* 1 (SI Fig. 8).

We also studied the effect of tertiary structure by looking at asymmetry in surface accessibility. *β* sheets at the N terminal are less likely to be exposed to solvent than *β* sheets at the C terminal; this bias is stronger for prokaryotic proteins (first 20 residues: 41 %) compared to eukaryotic proteins (13 %) (SI Fig 9). Since solvent-exposed *β*-sheets are less likely to form part of a folding nucleus [73], this suggests that *β*-sheets at the N terminal are more likely to nucleate folding compared to those at the C terminal.

### Correlations support the ‘slowest-first’ hypothesis

The data used to fit Eq. 7 are sparse (122 proteins), biased towards small, single-domain proteins, and typically obtained from *in vitro* refolding experiments [45]. To test whether our conclusions are robust to sampling, we estimate confidence intervals using bootstrapping with sample sizes equal to the original sample size, and half that amount; we perform this test on both the reduced version of the PFDB data set used in the main figures, and on a second version of the PFDB data set (Methods; SI Fig. 4). In addition, we calculate the main results using using a different protein folding data set, ACPro [74], which partially overlaps with PFDB, but includes larger proteins (SI Fig. 10). In all of the above analyses, the point of maximum asymmetry is found to be 1*/*100 *< R <* 100, which corresponds to the region where CTF speed-up is possible. However, to fully overcome the aforementioned limitations, further experiments are needed.

### Analysis is consistent with hypothesis that proteins are selected for CTF via secondary structure

To sum, in the proposed the *slowest-first* mechanism, CTF can be accelerated by positioning the slowest-folding parts of a protein near the N terminal so that they have more time to fold. A survey of the PDB shows that the estimated acceleration correlates with asymmetry in secondary structure. In particular, the rate of production can be almost doubled when translation time is similar to folding time, and indeed these proteins exhibit the maximal asymmetry in secondary structure distribution. Altogether, there appears to be substantial evolutionary selection, manifested in sequence asymmetry, for proteins that can fold during translation.

## Methods

### Data

We extracted a set of 16,200 proteins from the Protein Data Bank (PDB) [40]. We only include proteins that exactly match their Uniprot sequence (not mutated, spliced, or truncated) [75]. For each unique protein sequence, we only include the most recent structure. We used SIFTS to map PDB and Uniprot entries [76]. We exclude proteins with predicted signal peptides as little is known about whether such proteins undergo CTF; we used Signal-P5.0 to identify signal peptides [77]. Using the above criteria we extracted a set of 38,274 domains by matching PDB entries to Pfam domains [54]. *α* helix and *β*-sheets are identified through annotations in the PDB; disorder is inferred from residues with missing coordinates. To calculate contact order, we only consider contacts between residues where *α* carbons are within 10 Å; we confirm that the correlation in Fig. 1D is robust to choice of this cutoff (SI Fig. 11).

We use the protein folding kinetics database (PFDB) for estimating folding rates [45]. For our main results we only used entries with realistic physical conditions (5 *<* pH *<* 8, and 20 *°*C *< T <* 40 *°*C) and ignored folding rates which had been extrapolated to *T* = 25 *°*C; in total, 122 proteins. We test a second version of the PFDB data set without excluding proteins, and using folding rates which were extrapolated to *T* = 25 *°*C; 141 proteins. We also use the ACPro data set [74] to test the robustness of our conclusions; 125 proteins.

### Predicting folding and translation rate

The folding rate, *k*_fold_ (in units of 1/sec), is estimated by a power-law fit as a function of the protein’s length:

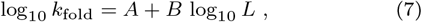

where *L* is sequence length in residues; *A* and *B* are free parameters. We fit these parameters using data from the PFDB [45] to get 95 % confidence intervals of *A* = 13.8 *±* 2.1 and *B* = *−*6.1 *±* 1.2 (with correlation coefficient *r* = *−*0.68, and p-value *p <* 0.005). The estimate from Eq. 7 is limited for the following reasons: (i) It is extracted from a small set of 122 proteins. (ii) It disregards the effects of secondary structure, contact order order, and other important determinants. (iii) The data is from *in vitro* measurements. (vi) The data is biased towards small, single-domain proteins. Thus, it is only a rough predictor for the folding rates of *individual* proteins in the set, as the standard deviation between estimated and empirical folding rates is 1.22. For all these reasons, we use Eq. 7 as an estimator of the *average* folding rate of sets of proteins of similar length *L* where the large sampling size of each bin is expected to reduce the errors as *∼ N*^*−*1*/*2^.

We tested whether the predicted folding rates of proteins in the PDB are within certain approximate bounds on realistic folding rates. A lower bound to folding time has been estimated at *∼ L/*100 µs [78], while we take the doubling time of *E. coli*, roughly 20 minutes, as an approximate upper bound. Of course, many proteins rely on chaperones, so their bare estimated folding time may be longer than the upper bound, while others come from organisms with much longer doubling times. Even so, according to Eq. 7 only 8 % of proteins are estimated to have a folding time greater than 20 minutes, while only 7 % of proteins are estimated to fold faster than the lower bound. Given the magnitude of the error in estimating the folding time of individual proteins, Eq. 7 appears to yield estimates that are mostly within the biologically reasonable regime. Furthermore, in estimating the folding rate of large proteins, a common assumption is that they consist of multiple independently-folding domains [79] – which considerably reduces the estimated folding time of the slowest proteins – but we neglect to make this assumption.

In principle, we could have used structural/topological measures (such as contact order, long-range order, etc. [80]) to slightly improve the fit to Eq. 7. However these typically involve numerous methodological choices and additional parameters [81], and the scaling relations are entirely empirical. In contrast, scaling of folding time with length has a robust theoretical background [44, 82–88]; the exact form of of the scaling is debated, but a power law is favoured slightly [83, 89].

We assume the translation rate, *k*_trans_, depends on the organism (host organism for viral proteins), such that *k*_trans_ is 5 amino acids per second for eukaryotes and 10 for prokaryotes.

## Data Availability

The non-redundant sets of proteins and domains, along with the data used in the figures and Supplementary Information, will be made available on Zenodo.

## Code Availability

Simulation and analysis code, along with code used to make all figures, are accessible a t https://github.com/jomimc/FoldAsymCode.

### Acknowledgements

We acknowledge Albert J. Libchaber for stimulating discussions and comments on the manuscript. This work was supported by the Institute for Basic Science, Project Code IBS-R020-D1.

## Author Contributions

J.M. and T.T. designed research; J.M. performed research; J.M. analyzed data; J.M. and T.T. wrote the paper.

## 0.1 Coarse-grained simulation of co-translational folding

In this work, we assume that *β*-sheets fold slower than *α*-helices in general, and hence our model predicts that co-translational folding (CTF) will finish faster if *β*-sheets are located at the N terminal. Here we further examine this phenomenon using coarse-grained models of proteins, such that we also take into account the effect of tertiary structure. We study three proteins that exhibit significant asymmetry in secondary structure, with native structures taken from the Protein Data Bank (PDB): 1ILO, 2OT2, 3BID.

To investigate whether these proteins benefit from having β-sheets at the N terminal, we first calculate the time it takes to fold after starting from an unfolded configuration, τ_fold_. Then, for a range of translation times (*τ*_trans_ = *Lτ*_AA_, where *τ*_AA_ is the time it takes to translate one amino acid) we calculate the time it takes to undergo translation and folding, *τ*_CTF_, for both translation from the N- to the C-terminal (*τ*_CTF,N_), and also from the C- to the N-terminal (*τ*_CTF,C_). We then plot the speed-up, the ratio between the time it takes for CTF along the backward to the forward direction (*τ*_CTF,C_*/τ*_CTF,N_), against the ratio of folding translation time, *R* = *τ*_fold_*/τ*_trans_ (Fig. 8). All three proteins folded faster (up to 21%) when *β*-sheets were translated first, and the maximum speed-up occurred when 1*/*10 *< R <* 10.

We run molecular-dynamics simulations using the Gromacs software package [1]. We use a Go-like model forcefield that represents each residue with a single bead, generated using the Go-Kit python package [2]. To calculate folding time, *τ*_fold_, we run simulations from the unfolded state at *T* ^*\**^ = 100, with a time step of *dt*^*\**^ = 0.001 for 2 *×* 10^6^ steps; units are reported in reduced units according to the definitions given by Gromacs. We record folding time, *τ*_fold_, as the time it takes for 90% of native contacts to be within 20% of the distance given in the PDB structure. We run the simulation 1,000 times to get an average.

To calculate co-translational folding time, *τ*_CTF,N_ (*τ*_CTF,C_), we initialise the protein in an elongated configuration and restrain all beads except the first bead at the N (C) terminal. We run the simulation for *τ*_AA_, the time it takes to translate a single residue, and then remove the restraint from the next bead in the chain; we repeat until all restraints are removed. We also create wall out of purely repulsive spheres packed on a square lattice, positioned at the boundary between free and restrained beads, which prevents interaction between the ‘translated’ and ‘untranslated’ parts of the protein. As each bead has its restraint removed, we move the wall along the chain. When the restraint is removed from the final bead, we remove the wall and run for a further 5 *×* 10^5^ or 1 *×* 10^6^ time steps. We measure *τ*_CTF,N_ (*τ*_CTF,C_) in the same way as *τ*_fold_. We run the simulation in each direction 4,000 times to get an average. A full set of simulation parameters is available in the supplementary materials in the form of Gromacs input files and python scripts for configuration set-up and analysis.

**Figure 1:**
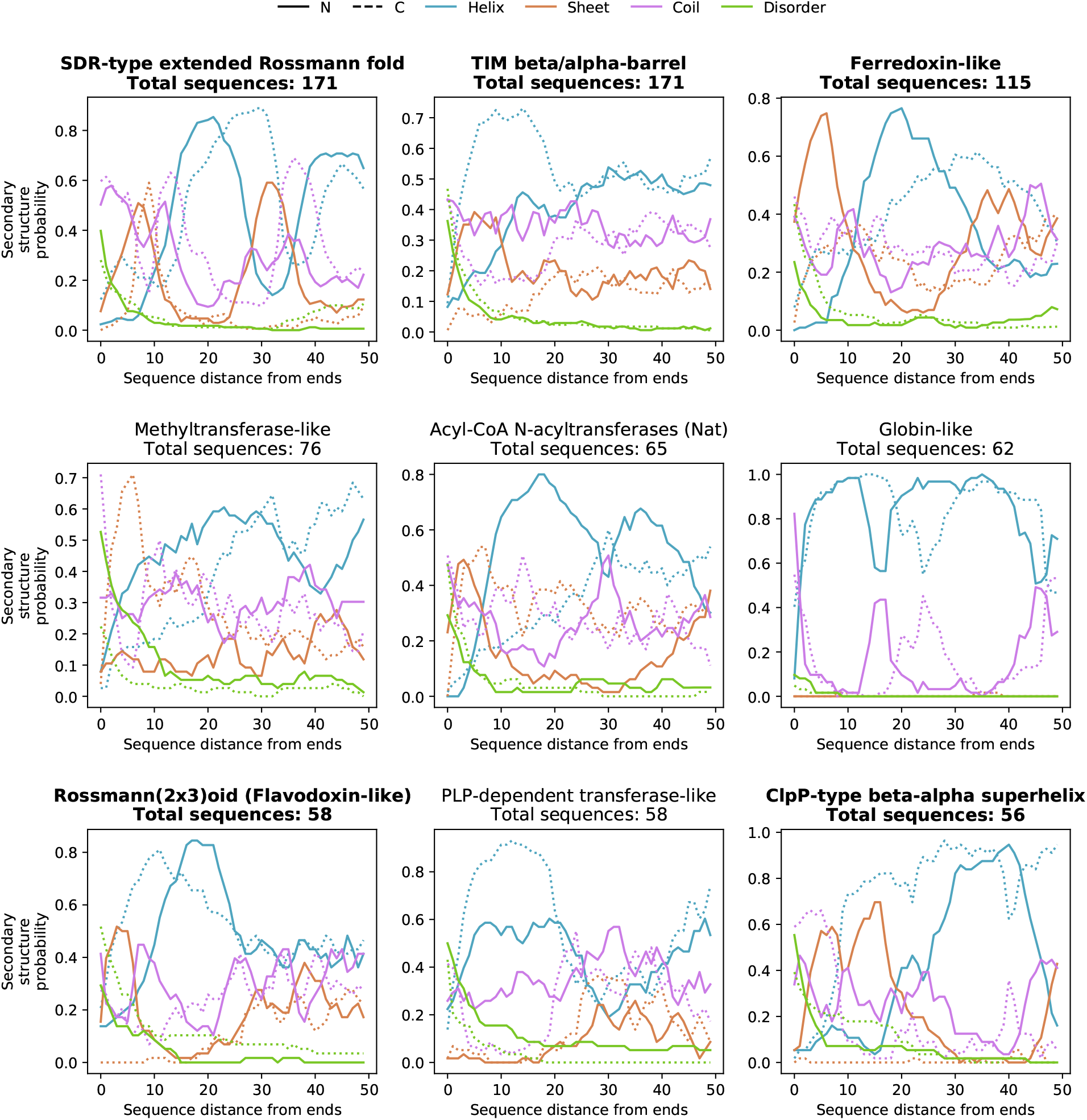
Secondary structure probability as a function of sequence distance from the N and C terminal respectively, for *α*-helices, *β*-sheets, random coil, and disorder. Separate plots are shown for the 9 most common SCOP folds in our sample of PDB proteins [3]. The folds that exhibit *α*-*β* periodicity in their secondary structure are highlighted in bold.

**Figure 2:**
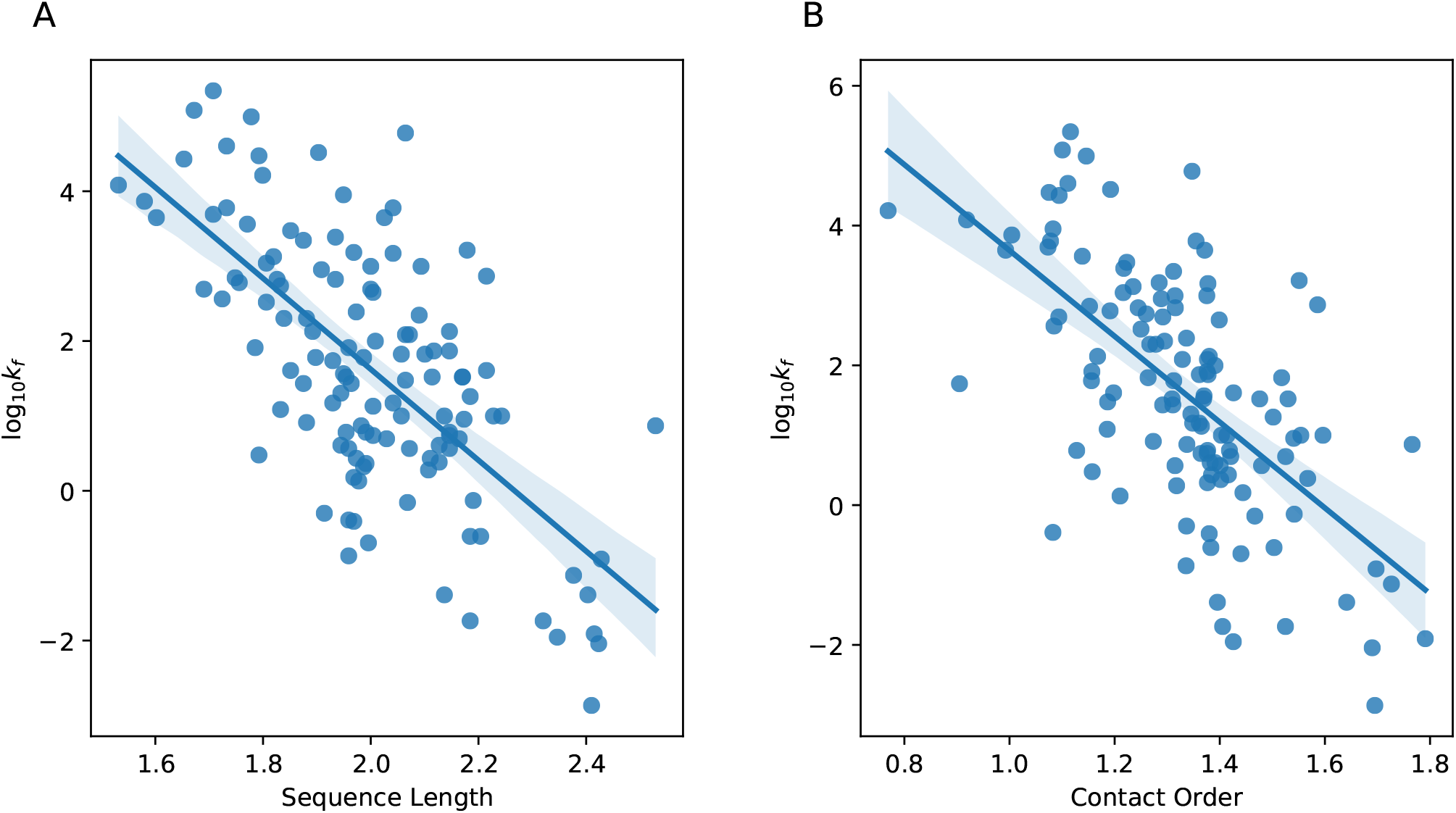
Correlations between sequence length (**A**), contact order (**B**) and empirically-determined folding rate *k*_*f*_ [4]. Two tailed Pearson’s correlation gives: sequence length, *r* = *−*0.68, *p <* 0.005; contact order, *r* = *−*0.71, *p <* 0.005.

**Figure 3:**
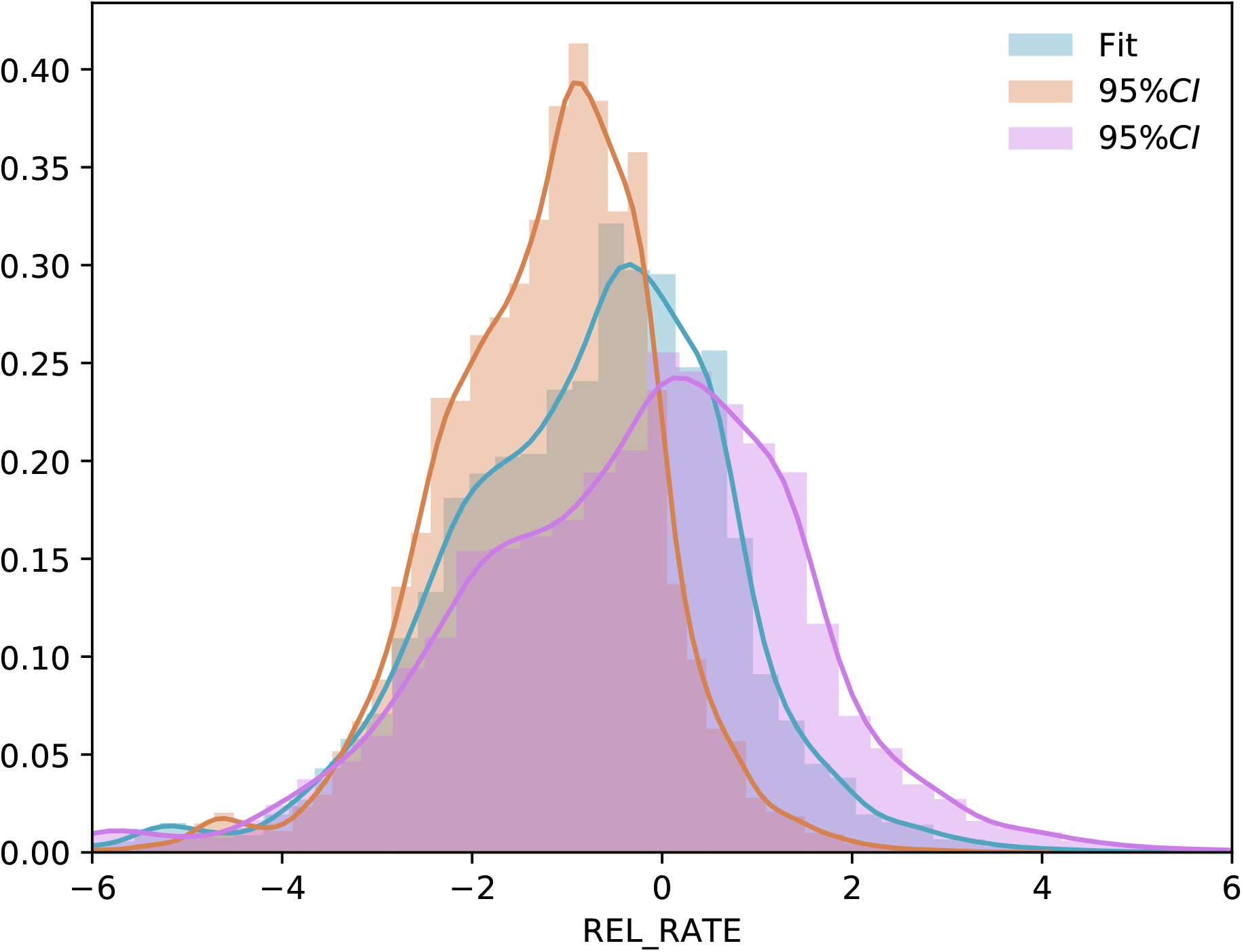
Distribution of *R*, the ratio of folding to translation time, estimated by using the main fit to Eq. 7 (main text), and the 95% confidence intervals to the fit.

**Figure 4:**
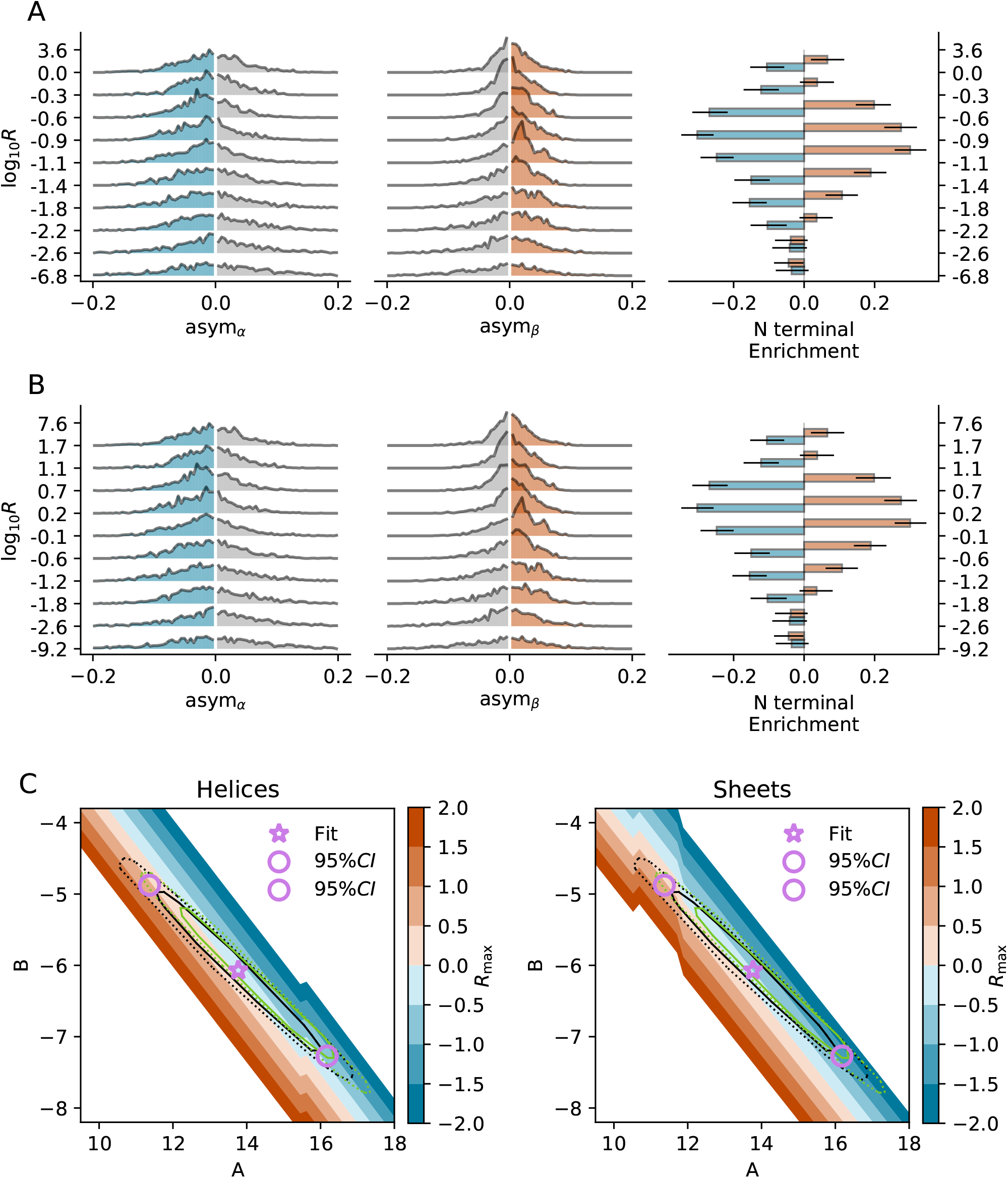
**A**-**B**: *α*-*β* asymmetry distributions and N terminal enrichment for the full sample according to the 95% confidence intervals on the empirical fit for estimating *k*_trans_. Proteins are divided into deciles according to *R*; bin edges are shown on the x-axis. **C**: The median of the *R* decile that exhibits maximum asymmetry, *R*_max_, as a function of fitting parameters *A* and *B*. The pink star corresponds to the fit used in the main text. The pink circles correspond to the fits used in **A** and **B**. The rings show the bootstrapped 95% confidence interval for values of *A* and *B*. Black rings are for the PFDB data set used in the main figures; green rings are for the less conservative PFDB data set. Solid rings indicate bootstrapped samples with the same sample size as the original sample; dotted rings indicate bootstrapped samples with half the sample size of the original sample.

**Figure 5:**
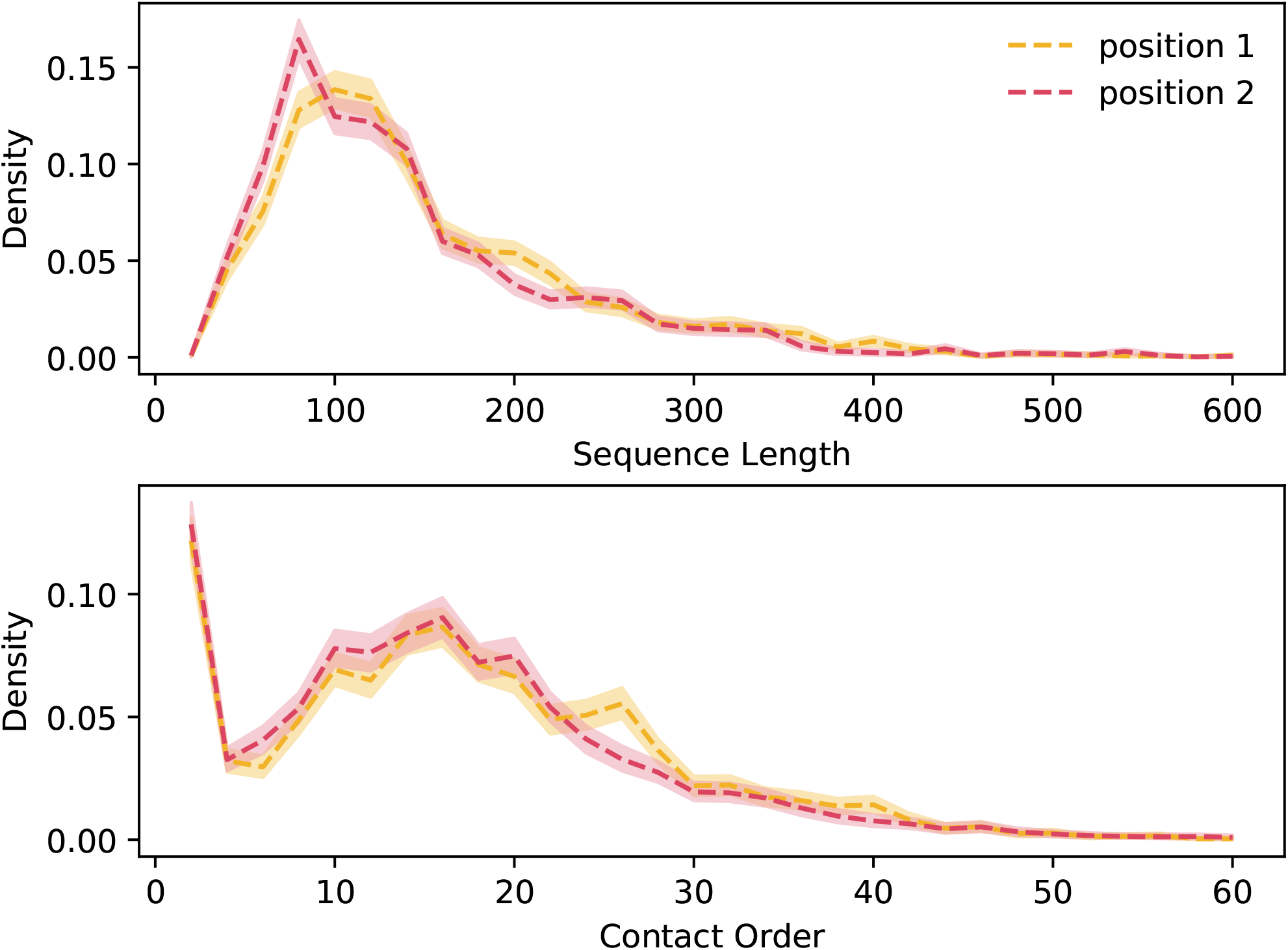
Sequence length (top) and contact order (bottom) distributions for domains in position 1 (near the N terminal) and position 2 in two-domain proteins. Shaded area indicates bootstrapped 95% confidence intervals. The distributions do not differ based on domain position, which suggests that folding time does not depend on domain position.

**Figure 6:**
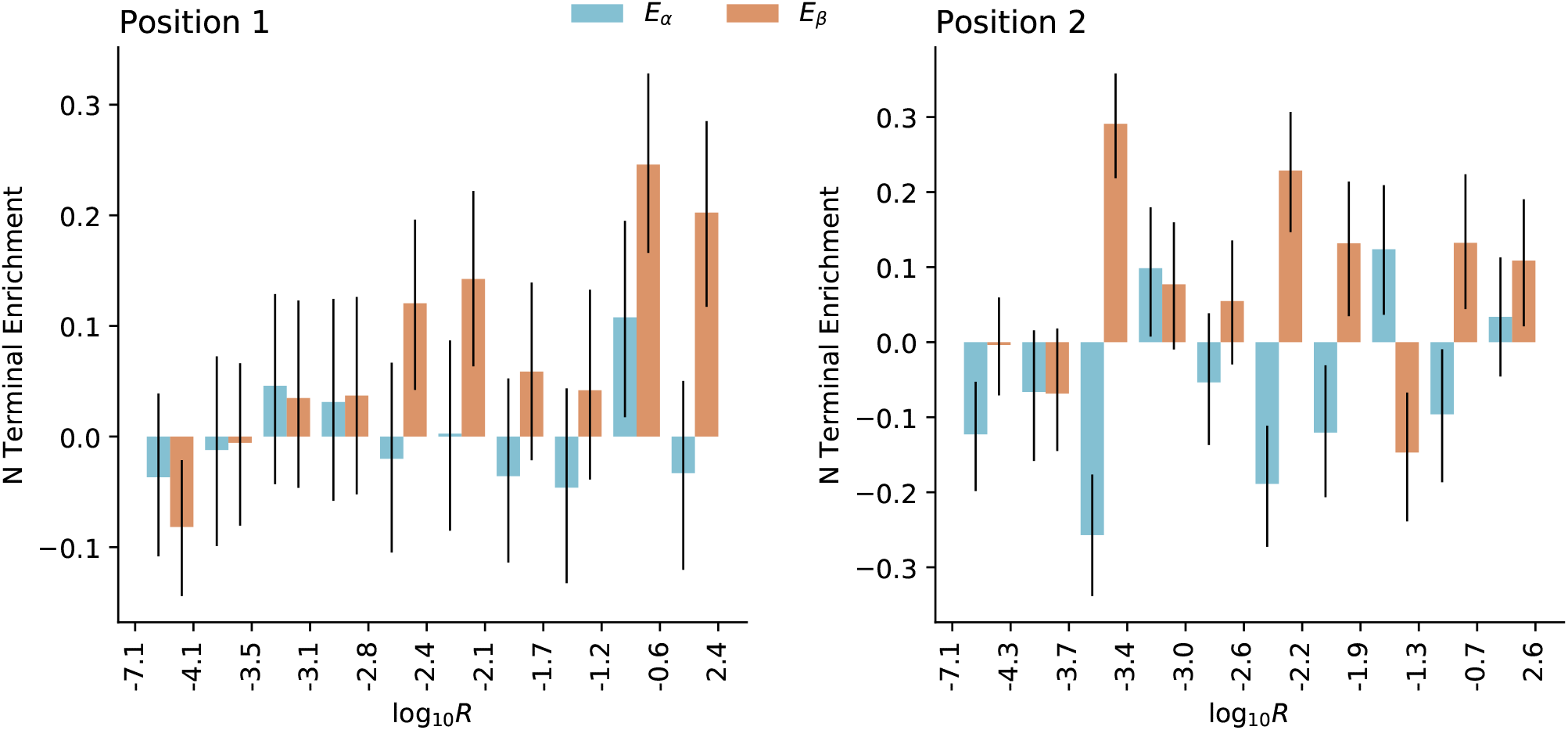
N terminal enrichment of *α* helices (*E*_*α*_) and *β* sheets (*E*_*β*_) for individual domains within two-domain proteins for domains in position 1 (left; closest to the N terminal) and position 2 (right). Proteins are divided into deciles according to *R*; bin edges are shown on the x-axis; whiskers indicate bootstrapped 95% confidence intervals. The data shows that domains in position 1 exhibit maximum *β* asymmetry when *−*1.2 *< R <* 2.4, in agreement with the proposed ‘slowest-first’ scheme. However, the confidence intervals are almost as large as the effect size, so we lack sufficient data to show a significant effect. Domains in position 2 do not appear to exhibit asymmetry in agreement with the ‘slowest-first’ scheme.

**Figure 7:**
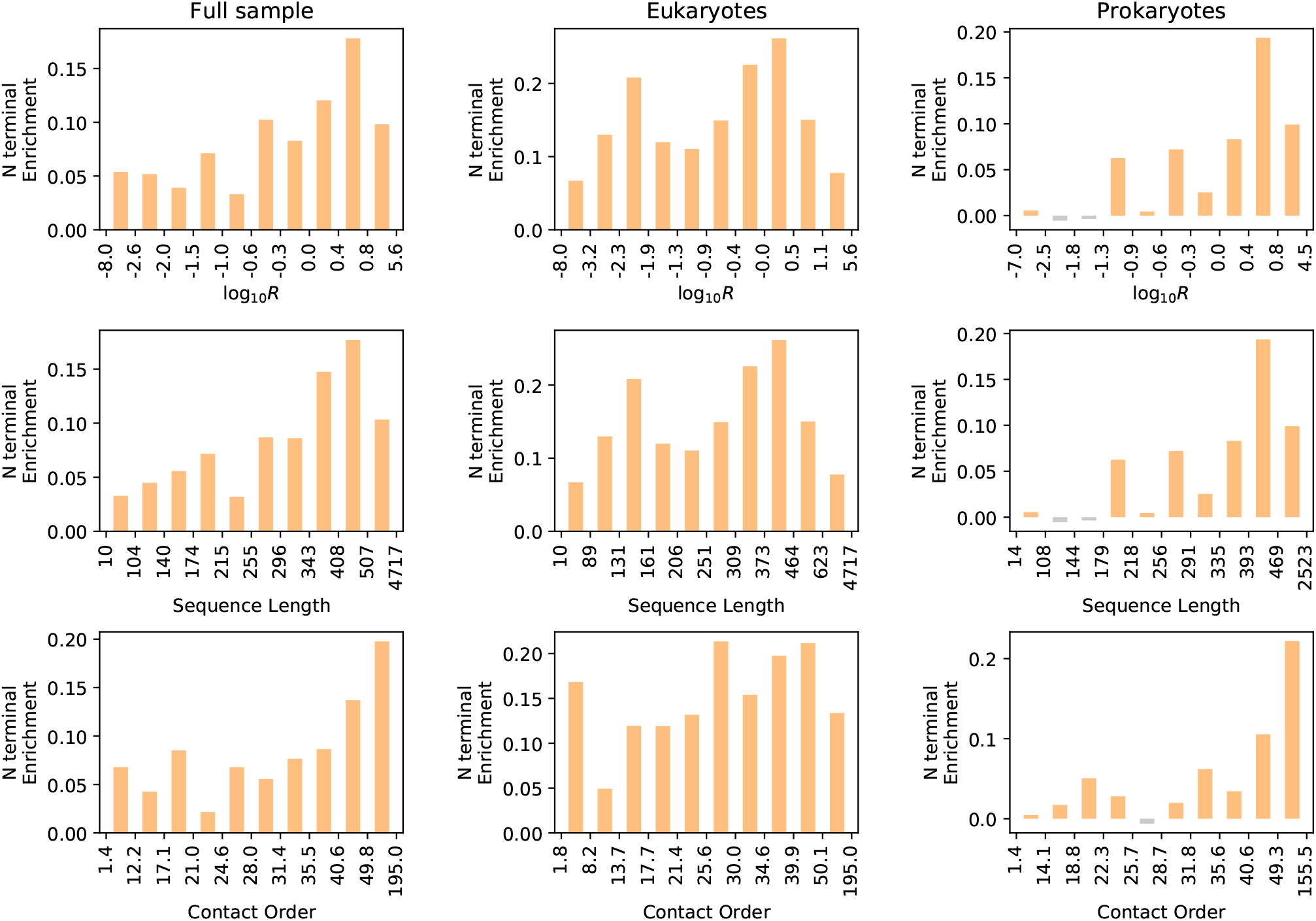
N terminal enrichment of disordered residues as a function of log_10_ *R*, Sequence Length, and Contact Order; data is shown for the full sample (left), eukaryotic proteins (middle), and prokaryotic proteins (right). Proteins are divided into deciles according to *R* (top), Sequence Length (middle), and Contact Order (bottom); bin edges are shown on the x-axis. There is a clear association between N terminal enrichment and slow-folding proteins in prokaryotes, but not in eukaryotes.

**Figure 8:**
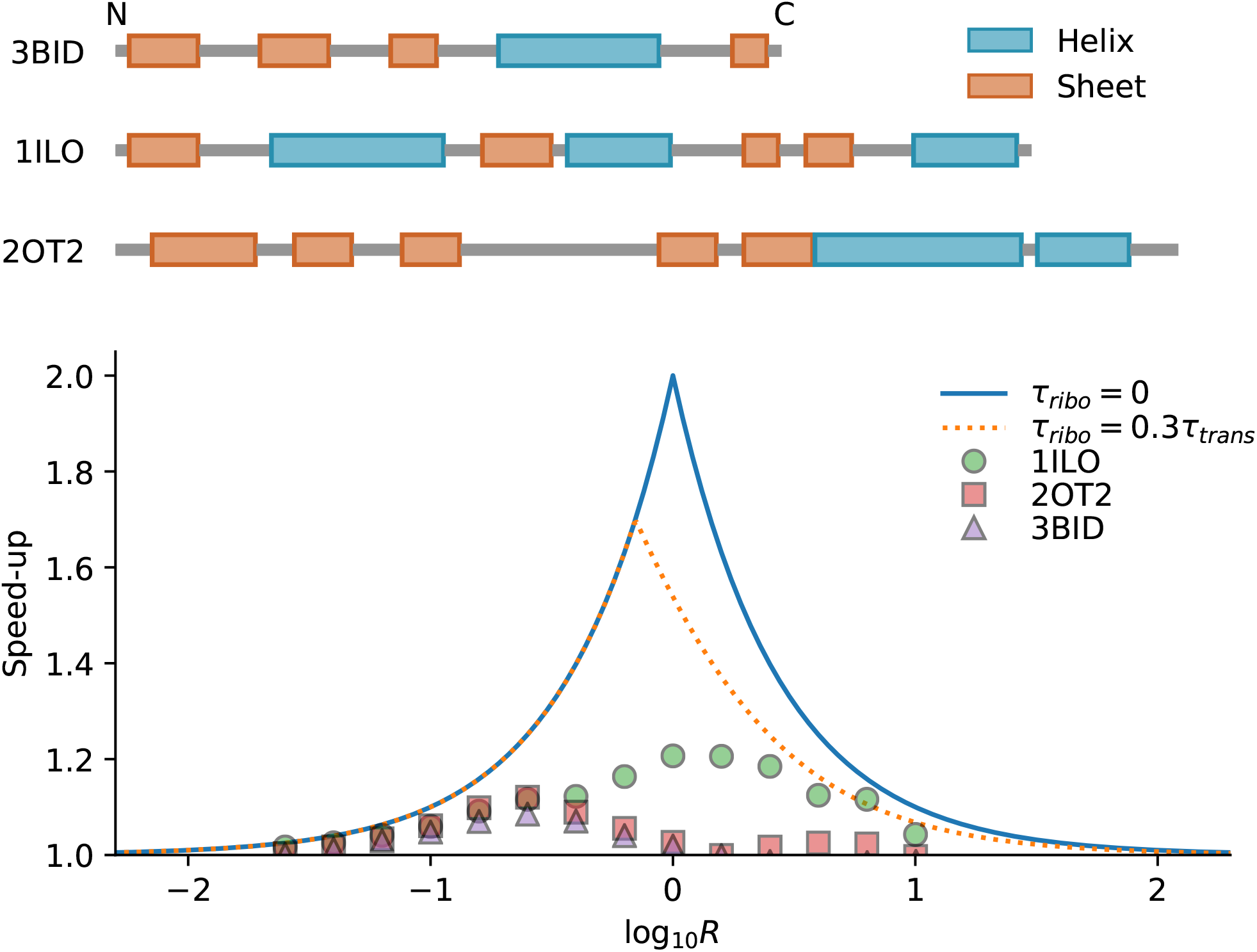
(Top) Secondary structure of three proteins that exhibit *α*-*β* asymmetry in line with the ‘slowest-first’ scheme (PDB IDs: 3BID, 1ILD, 2OT2). (Bottom) Maximum theoretical speed-up achievable as a function of *R* and *τ*_ribo_ (lines). Speed-up achieved in a simulation by translating proteins from the N- to the C-terminal (with slow-folding parts near the N terminal) compared to translating them from the C- to the N-terminal.

**Figure 9:**
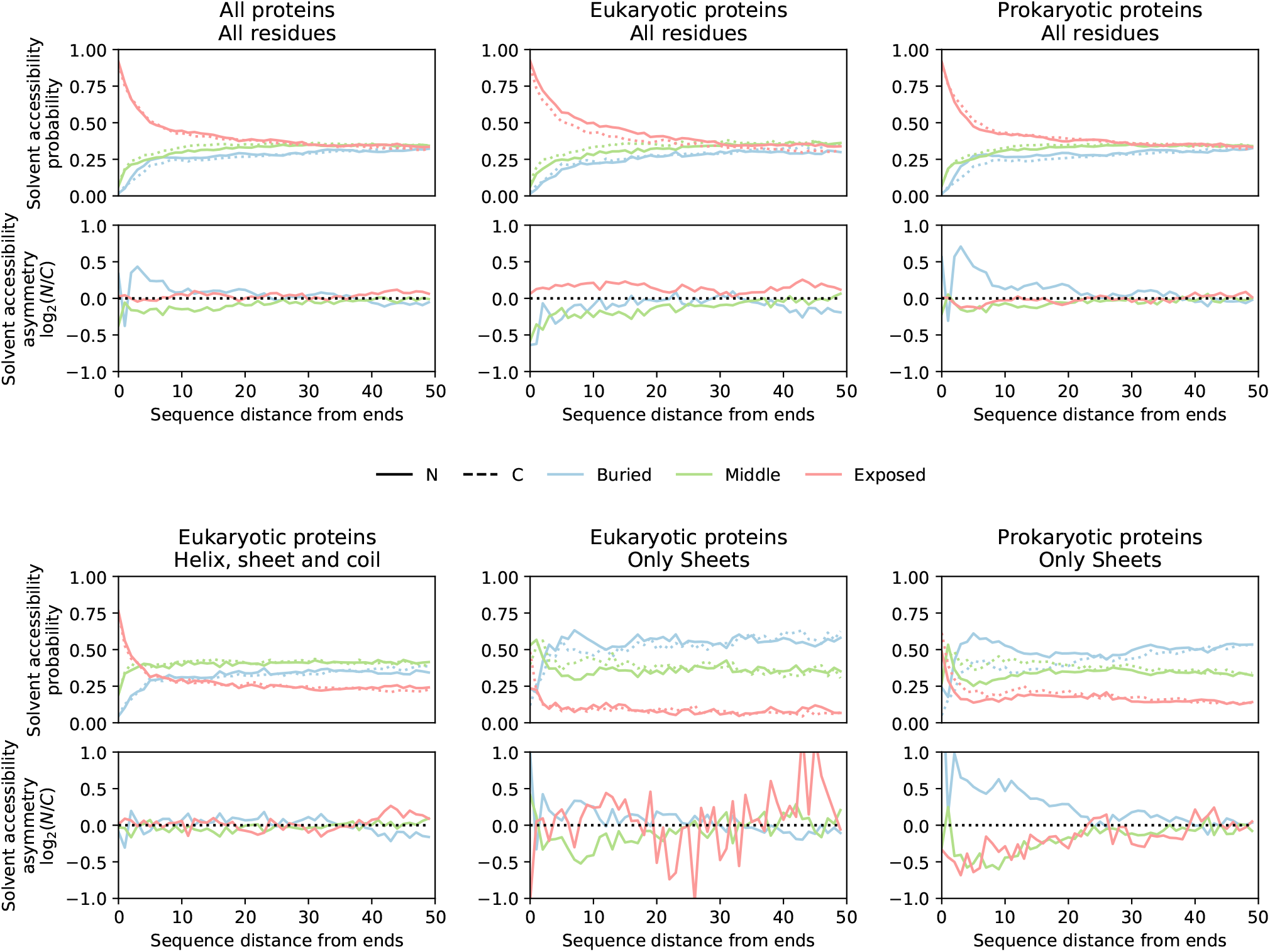
Relative solvent accessibility (SA) and SA asymmetry as a function of sequence distance from the terminal residues. SA is calculated using the freesasa C library [5, 6], and residues are divided equally among three categories: Buried, Middle, or Exposed. It is impossible to calculate SA for disordered residues, and we assume that these are Exposed [7]. (Top) Data is shown for all proteins (left), eukayotic proteins (middle) and prokaryotic proteins (right). (Bottom) Data is shown for helix, sheet and coil residues from eukaryotic proteins (left), for sheet residues from eukarotic proteins (middle), and for sheet residues from prokaryotic proteins (right). For eukaryotic proteins overall, the N terminal has a tendency to be more exposed than the C terminal. However, when only considering helices, sheets and coils there is no significant asymmetry – *i*.*e*. the bias for being exposed at the N terminal is explained by the bias for being disordered at the N terminal. When only considering sheets, the N terminal is slightly more buried, while the C terminal is more likely to contain ‘Middle’ values of SA. For prokaryotic proteins overall, the N terminal is more likely to buried compared to the C terminal, which is mainly due to *β* sheets.

**Figure 10:**
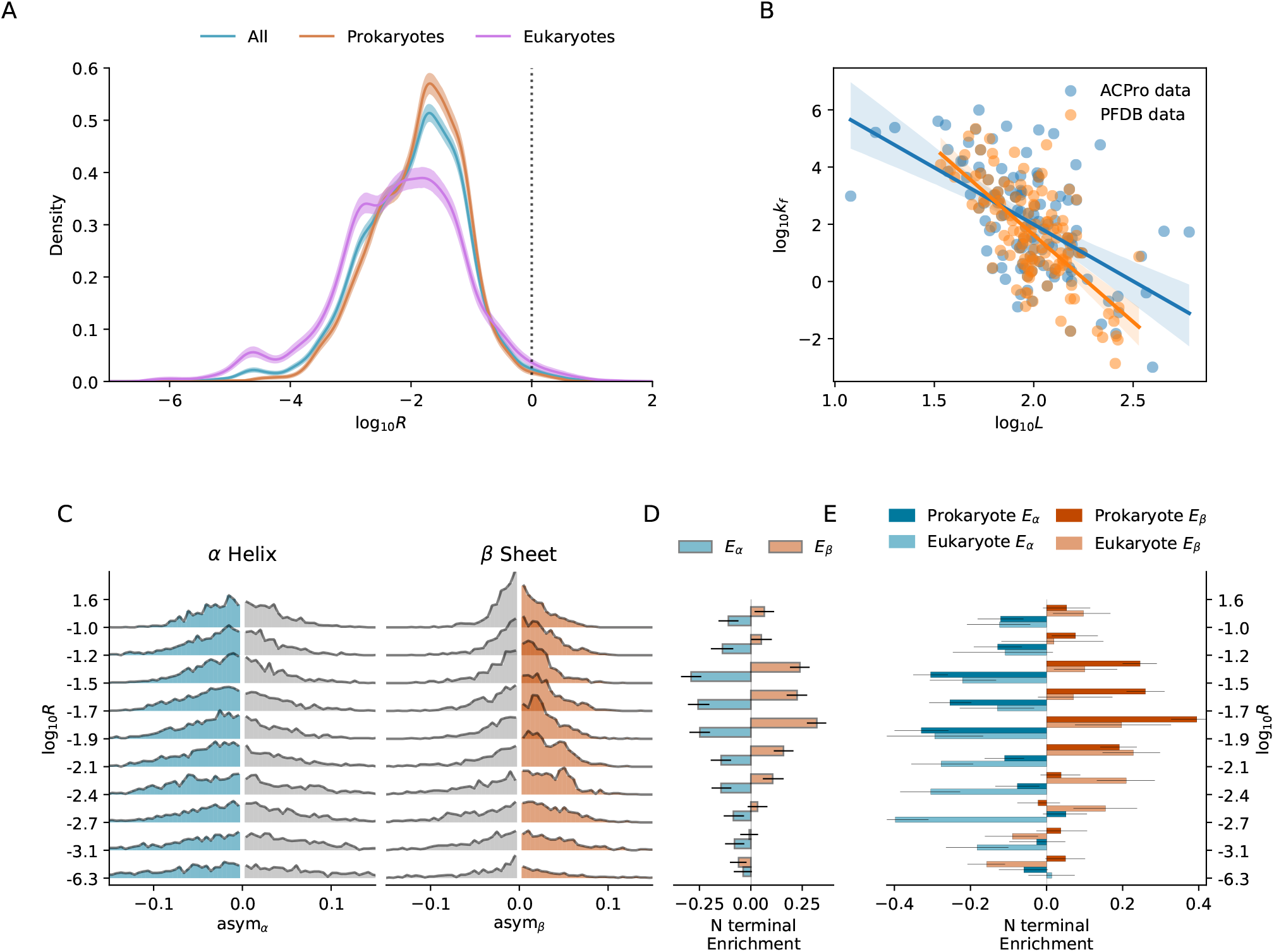
Repeat of main text Figures 2-3 after calculating *R*, the folding/translation time ratio, using the ACPro database [8] instead of the PFDB [4]. **A**: *R* distribution for the full PDB sample, prokaryotic proteins, and eukaryotic proteins. **B**: Scatter plot and correlation between sequence length *L* and folding rate *k*_*f*_ for both the ACPro and PFDB databases. Shaded area indicates 95% confidence interval of the linear fit. **C**: *α*-*β* asymmetry distributions as a function of *R*. Proteins are divided into deciles according to *R*; bin edges are shown on the y-axis. **D**: N terminal enrichment – the degree to which sheets/helices are enriched in the N over the C terminal – is shown for the deciles given in C. **E**: N terminal enrichment as a function of *R* for 4,633 eukaryotic proteins and 10,400 prokaryotic proteins. Proteins are divided into bins according to *R*; bin edges, shown on the x-axis, are the same as in C-D. Whiskers indicate 95% confidence intervals. In contrast with the results obtained using the PFDB, estimating *R* using the ACPro database results in the prediction that for most proteins translation time is considerably longer than folding time. The principal reason for this appears to be a few proteins with either few residues (*<* 34) or many residues (*>* 300) that are present in the ACPro data but not the PFDB. These residues change the gradient of the fit such that large proteins are predicted to fold much faster according to the ACPro fit. Manavalan et al. [4] cite several reasons for why certain entries in ACPro were not used in the PFDB. Even despite these differences, the region of maximal asymmetry is predicted to be *−*1.9 *< R < −*1.2, where non-negligible acceleration of CTF is still possible.

**Figure 11:**
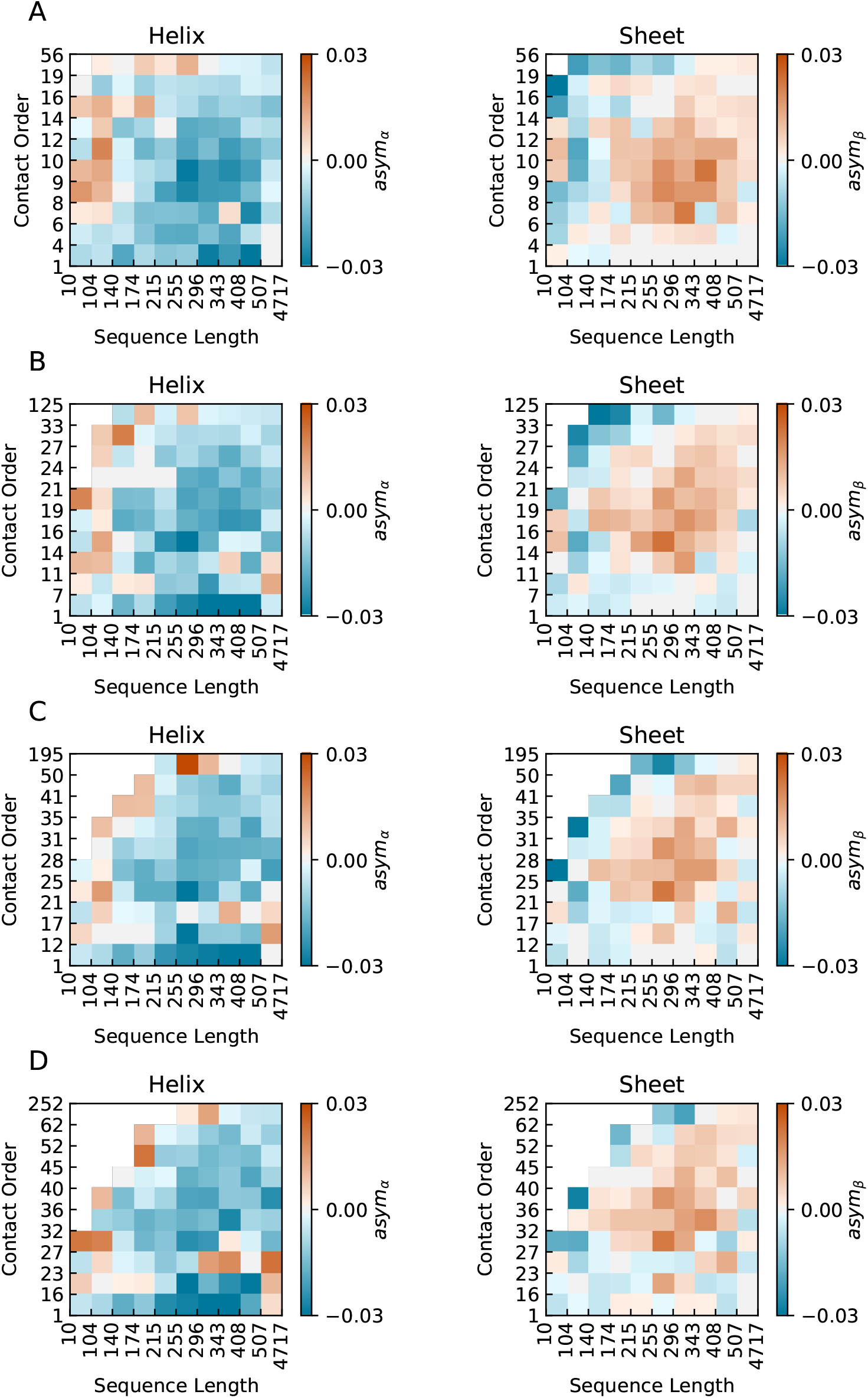
Correlation between sequence length, contact order and *α*-*β* asymmetry. Separate plots are shown for different values of the cutoff used to calculate contact order: **A**, 6 Å; **B**, 8 Å; **C**, 10 Å; **D**, 12 Å.

